# Parallel paleogenomic transects reveal complex genetic history of early European farmers

**DOI:** 10.1101/114488

**Authors:** Mark Lipson, Anna Szécsényi-Nagy, Swapan Mallick, Annamária Pósa, Balázs Stégmár, Victoria Keerl, Nadin Rohland, Kristin Stewardson, Matthew Ferry, Megan Michel, Jonas Oppenheimer, Nasreen Broomandkhoshbacht, Eadaoin Harney, Susanne Nordenfelt, Bastien Llamas, Balázs Gusztáv Mende, Kitti Köhler, Krisztián Oross, Mária Bondár, Tibor Marton, Anett Osztás, János Jakucs, Tibor Paluch, Ferenc Horváth, Piroska Csengeri, Judit Koós, Katalin Sebők, Alexandra Anders, Pál Raczky, Judit Regenye, Judit P. Barna, Szilvia Fábián, Gábor Serlegi, Zoltán Toldi, Emese Gyöngyvér Nagy, János Dani, Erika Molnár, György Pálfi, László Márk, Béla Melegh, Zsolt Bánfai, László Domboróczki, Javier Fernández-Eraso, José Antonio Mujika-Alustiza, Carmen Alonso Fernández, Javier Jiménez Echevarría, Ruth Bollongino, Jörg Orschiedt, Kerstin Schierhold, Harald Meller, Alan Cooper, Joachim Burger, Eszter Bánffy, Kurt W. Alt, Carles Lalueza-Fox, Wolfgang Haak, David Reich

## Abstract

Ancient DNA studies have established that European Neolithic populations were descended from Anatolian migrants who received a limited amount of admixture from resident hunter-gatherers. Many open questions remain, however, about the spatial and temporal dynamics of population interactions and admixture during the Neolithic period. Using the highest-resolution genome-wide ancient DNA data set assembled to date—a total of 177 samples, 127 newly reported here, from the Neolithic and Chalcolithic of Hungary (6000–2900 BCE, *n* = 98), Germany (5500–3000 BCE, *n* = 42), and Spain (5500–2200 BCE, *n* = 37)—we investigate the population dynamics of Neolithization across Europe. We find that genetic diversity was shaped predominantly by local processes, with varied sources and proportions of hunter-gatherer ances try among the three regions and through time. Admixture between groups with different ancestry profiles was pervasive and resulted in observable population transformation across almost all cultural transitions. Our results shed new light on the ways that gene flow reshaped European populations throughout the Neolithic period and demonstrate the potential of time-series-based sampling and modeling approaches to elucidate multiple dimensions of historical population interactions.

---

The population dynamics of the Neolithization process are of great importance for understanding European prehistory [1–5]. The first quantitative model of the Neolithic transition to integrate archaeological and genetic data was the demic diffusion hypothesis [1], which posited that growing population densities among Near Eastern farmers led to a range expansion that spread agriculture to Europe. Ancient DNA analysis has validated major migrations from populations related to Neolithic Anatolians as driving the arrival of farming in Europe [6–13], but the demic diffusion model does not account for the complexities of the interactions between farmers and hunter-gatherers in Europe throughout the Neolithic [2,3,14–16]. For example, ancient DNA has shown that farmers traversed large portions of Europe with limited initial admixture from hunter-gatherers [8,10,12,13,17], and furthermore that farmers and hunter-gatherers lived in close proximity in some locations long after the arrival of agriculture [18, 19]. However, genetic data have yet to be used systematically to model the population interactions and transformations during the course of the Neolithic period. Key open questions include whether migrating farmers mixed with hunter-gatherers at each stage of the expansion, and if so how soon after arriving. Additionally, while it has previously been shown that hunter-gatherer ancestry among farmers in several parts of Europe had increased by the Middle Neolithic [10, 12, 13, 20], it is currently unclear whether this was a continuous or discrete process and furthermore whether it involved a continent-wide phenomenon or a variety of parallel, local events.

We compiled a high-resolution data set of 177 Neolithic and Chalcolithic European genomes (pre-dating the arrival of steppe ancestry in the third millennium BCE [10]) from what are now Hungary, Germany, and Spain, of which 127 individuals are newly reported here, 38 with new direct radiocarbon dates (Figure 1A, B; Extended Data Table 1; Supplementary Tables 1, 2; Supplementary Information sections 1–3). We enriched for DNA fragments covering a set of ∼1.23 million single nucleotide polymorphism (SNP) targets [12, 21] (Methods) and obtained largely high-quality data, with at least 100,000 SNPs hit at least once (average coverage ∼0.1 or higher) for 88 of the 127 samples. The majority (88) of our new samples comprise an approximately 3000-year transect of the prehistory of the Carpathian Basin (Supplementary Information section 1), from both the eastern (Great Hungarian Plain, or Alföld) and western (Transdanubia) portions of present-day Hungary: 7 Early Neolithic (EN, ∼6000–5500 BCE) Starčevo (western) and Körös (eastern) individuals; 33 Middle Neolithic (MN, ∼5500–5000 BCE) individuals from the Transdanubian (LBKT) and Alföld (ALPc) Linearbandkeramik cultures and from the Vinča culture in southern Transdanubia; 20 Late Neolithic (LN, ∼5000–4500 BCE) individuals (eastern Tisza, western Sopot and Lengyel, the latter two grouped together as Transdanubian LN, or TDLN); and 28 Tiszapolgár, Balaton-Lasinja, Hunyadihalom, Protoboleráz, and Baden individuals from the Chalcolithic period (CA, ∼4500–2850 BCE). From what is now Germany, we generated data for 15 new Linearbandkeramik (LBK) EN (∼5500–4850 BCE) and eight new MN (∼4600–3000 BCE) individuals (with new libraries for an additional nine LBK for which we have previously reported data), while from Spain, we sequenced two new EN (∼5500–4500 BCE) and 14 new CA (∼3000–2200 BCE) individuals. After quality control (Methods), we retained 110 samples, which we merged with 50 Neolithic individuals from the literature [9,10,12,22,23]. For population genetic analyses, we focused on a subset of 151 individuals from 15 population groupings for which we had the highest-quality data. We co-analyzed these samples with 25 Neolithic individuals (∼6500–6000 BCE) from northwestern Anatolia [12] to represent the ancestors of the first European farmers (FEF; Supplementary Information section 4) and four primary European hunter-gatherer individuals (“western hunter-gatherers,” or WHG): the ∼5700 BCE “KO1” from Hungary [12, 22], the ∼5900 BCE “La Braña 1” (LB1) from Spain [12, 24], the ∼6100 BCE “Loschbour” from Luxembourg [9], and the ∼12,000 BCE “Villabruna” from northeastern Italy [25].

**Figure 1.**
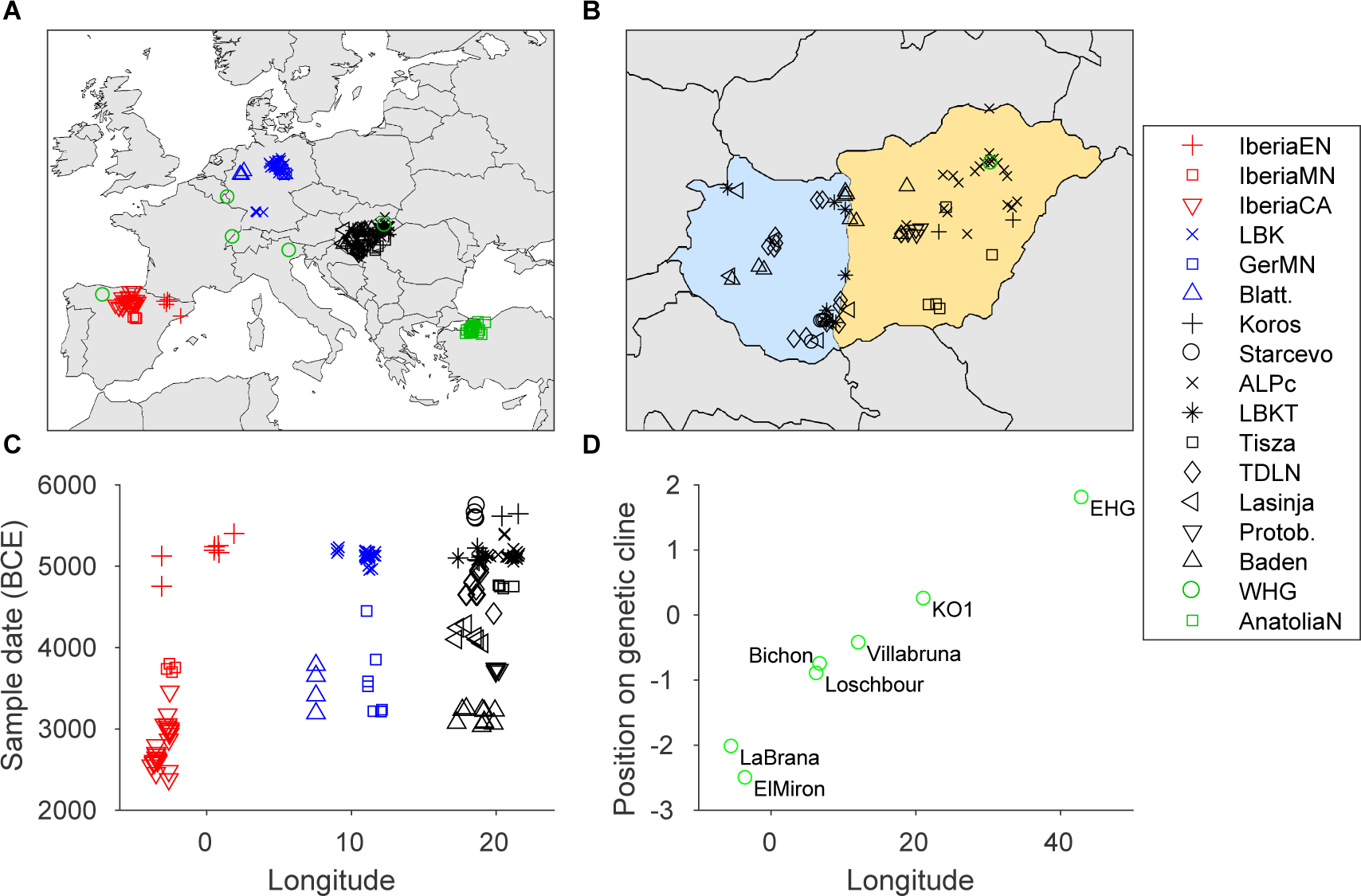
Spatial and temporal contexts of European Neolithic samples. (A), (B) Locations of samples used for analyses, with closeup of Hungary (orange shading for Alföld and light blue for Transdanubia). (C) Sample dates arranged by longitude. (D) Hunter-gatherer genetic cline (derived from MDS analysis; Supplementary Information section 5) as a function of longitude. Random jitter is added to separate overlapping positions in (A)–(C). GerMN, Germany MN; Blatt., Blätterhöhle; Protob., Protoboleráz.

**Extended Data Table 1.** Descriptive statistics, background information, and admixture parameters for Neolithic individuals in the study.

| ID | Population | Site | Lat. | Long. | Date | Sex | Mt Hap | Y Hap | Cov. | HG% | ALD | Ref. |
| --- | --- | --- | --- | --- | --- | --- | --- | --- | --- | --- | --- | --- |
| GEN68 | Körös | Törökszentmiklós road 4 site 3 | 47.2 | 20.4 | 5706–5541 | F | K1a | .. | 2.46 | -0.09±1.6 | ± |  |
| HUNG276, KO2 | Körös | Tiszaszőlős-Domaháza | 47.3 | 21.5 | 5713–5566 | F | K1a | .. | 0.91 | -1.51±1.6 | ± | [12, 22] |
| BAM17b | Starčevo | Alsónyék-Bátaszék, Mézőkerti telep | 46.2 | 18.7 | 5832–5667 | M | T1a2 | H2 | 1.47 | 7.85±1.7 | 4.5±1.9 |  |
| BAM25 | Starčevo | Alsónyék-Bátaszék, Mézőkerti telep | 46.2 | 18.7 | 5702–5536 | M | N1a1a1 | H2 | 0.22 | 1.63±1.9 | ± | [10, 12] |
| BAM4a | Starčevo | Alsónyék-Bátaszék, Mézőkerti telep | 46.2 | 18.7 | 5641–5547 | M | K1a4 | G2a2a1 | 0.20 | 3.43±2.0 | ± |  |
| LGCS1a | Starčevo | Lánycsók | 46.0 | 18.6 | 5800–5500 | M | W5 | G2a2b2b1a | 0.77 | -0.64±1.7 | ± |  |
| BAL25b | LBKT | Bátaszék-Lajvér | 46.2 | 18.7 | 5208–4948 | M | K1b1a | G2a2a1 | 2.77 | 0.06±1.5 | ± |  |
| BOVO1b | LBKT | Bölcske-Gyűrűsvölgy | 46.7 | 19.0 | 5300–4900 | F | H | .. | 0.01 | 11.1±6.4 | ± |  |
| BUD4a | LBKT | Budakeszi-Szőlőskert | 47.5 | 18.9 | 5300–4900 | M | T1a | G2a2b2a | 0.17 | 6.80±2.3 | 36±6.2 |  |
| BUD9a | LBKT | Budakeszi-Szőlőskert | 47.5 | 18.9 | 5300–4900 | F | U2 | .. | 1.10 | 1.89±1.6 | ± |  |
| GEN18 | LBKT | Alsónyék, site 11 | 46.2 | 18.7 | 5309–5074 | M | T2c1 | G2a2b2b1 | 1.48 | 2.69±1.5 | 35±12 |  |
| KON3 | LBKT | Enese elkerülő, Kóny, Proletár-dűlő, M85, Site 2 | 47.6 | 17.4 | 5300–4900 | F | T2b | .. | 0.03 | 2.82±4.1 | ± |  |
| SZEH4 | LBKT | Szemely-Hegyes | 46.4 | 18.7 | 5207–4944 | F | N1a1a1a3 | .. | 0.07 | 1.90±3.1 | ± | [10, 12] |
| CEG07b | ALPc | Cegléd, site 4/1 | 47.2 | 19.9 | 5300–4900 | M | J2b1 | G2a2b2a | 0.30 | 11.5±2.0 | ± |  |
| CEG08b | ALPc | Cegléd, site 4/1 | 47.2 | 19.9 | 5300–4900 | F | J1c1 | .. | 0.19 | 11.2±2.2 | 23±3.1 |  |
| EBSA2a | ALPc | Ebes-Sajtgár | 47.5 | 21.5 | 5300–4900 | F | K1a | .. | 0.05 | 16.4±3.1 | ± |  |
| EBVO5a | ALPc | Ebes-Zsongvölgy | 47.5 | 21.5 | 5300–4900 | M | V1a | CT | 0.04 | 9.36±3.3 | ± |  |
| HAE10a | ALPc | Hajdúnánás-Eszlári út | 47.9 | 21.4 | 5221–5000 | M | J2b1 | I | 0.29 | 11.0±1.8 | ± |  |
| HAE7a | ALPc | Hajdúnánás-Eszlári út | 47.9 | 21.4 | 5302–5057 | M | K1a | I2 | 1.57 | 9.26±1.8 | 6.2±5.7 |  |
| HELI11a | ALPc | Hejőkürt-Lidl | 47.9 | 21.0 | 5209–4912 | M | N1a1a1 | I2a2a1b | 0.99 | 6.08±1.8 | 14±2.0 |  |
| HELI2a | ALPc | Hejőkürt-Lidl | 47.9 | 21.0 | 5300–4900 | M | U8b1b | I | 0.09 | 7.48±2.6 | ± |  |
| HUNG302, NE2 | ALPc | Debrecen Tocopart Erdoalja | 47.5 | 21.6 | 5291–5056 | F | H | .. | 4.88 | 11.1±1.7 | ± | [12, 22] |
| HUNG372, NE5 | ALPc | Kompolt-Kígyóvár | 47.2 | 20.8 | 5295–4950 | M | J1c1 | C1a2 | 4.25 | 7.56±1.7 | ± | [12, 22] |
| HUNG86, NE3 | ALPc | Garadna-Elkerülő út site 2 | 48.5 | 21.2 | 5281–5026 | F | X2b-T226C | .. | 3.32 | 12.2±1.7 | 18±3.1 | [12, 22] |
| MEMO24b | ALPc | Mezőkövesd-Mocsolyás | 47.8 | 20.6 | 5500–5300 | M | U8b1b | CT | 0.04 | 11.8±3.3 | 26±12 |  |
| MEMO2b | ALPc | Mezőkövesd-Mocsolyás | 47.8 | 20.6 | 5500–5300 | F | K1a1 | .. | 2.28 | 9.10±1.7 | 24±5.2 |  |
| MEMO7a | ALPc | Mezőkövesd-Mocsolyás | 47.8 | 20.6 | 5500–5300 | F | HV | .. | 0.26 | 1.66±2.0 | 13±6.1 |  |
| PF325, NE1 | ALPc | Polgár-Ferenci-hát | 47.9 | 21.2 | 5306–5071 | F | U5b2c | .. | 1.52 | 8.21±1.8 | 11±3.9 | [12, 22] |
| PF839/1198, NE4 | ALPc | Polgár-Ferenci-hát | 47.9 | 21.2 | 5211–5011 | F | J1c5 | .. | 3.49 | 10.1±1.7 | 25±9.9 | [12, 22] |
| POPI5a | ALPc | Polgár-Piócás | 47.9 | 21.1 | 5300–4900 | M | K1a1 | I2a2a | 0.31 | 9.87±2.0 | 11±3.7 |  |
| PULE1.18a | ALPc | Pusztataskony-Ledence | 47.5 | 20.5 | 5300–4900 | F | T2c1d1 | .. | 0.29 | 10.8±1.9 | ± |  |
| PULE1.23a | ALPc | Pusztataskony-Ledence | 47.5 | 20.5 | 5300–4900 | F | H1e | .. | 0.17 | 9.63±2.2 | 11±3.4 |  |
| TISO13a | ALPc | Tiszadob-Ökenéz | 48.0 | 21.2 | 5208–4942 | M | J1c2 | I2a2a | 1.21 | 13.0±1.7 | 22±7.6 |  |
| TISO1b | ALPc | Tiszadob-Ökenéz | 48.0 | 21.2 | 5300–4900 | M | H7 | I2a2a1b1 | 0.11 | 7.32±2.5 | 29±13 |  |
| TISO3a | ALPc | Tiszadob-Ökenéz | 48.0 | 21.2 | 5300–4900 | F | U5b2b1a | .. | 0.27 | 12.3±2.1 | 8.4±5.2 |  |
| SEKU10a | Vinča | Szederkény-Kukorica-dűlő | 45.6 | 18.3 | 5320–5080 | M | K2a | G2a2b2a1a | 0.24 | 2.30±2.0 | ± |  |
| SEKU6a | Vinča | Szederkény-Kukorica-dűlő | 45.6 | 18.3 | 5321–5081 | F | H26 | .. | 1.15 | 9.27±1.7 | 9.1±9.4 |  |
| VEGI17a | Vinča | Versend-Gilencsa | 45.6 | 18.3 | 5400–5000 | F | U2 | .. | 0.01 | -6.22±5.6 | ± |  |
| VEGI3a | Vinča | Versend-Gilencsa | 45.6 | 18.3 | 5400–5000 | M | T2b | H2 | 0.41 | 0.54±1.8 | ± |  |
| Gorzs18 | Tisza | Hódmezővásárhely-Gorzsa | 46.4 | 20.4 | 5000–4500 | M | U5b2c | I2a1 | 6.87 | 7.86±1.7 | 13±4.2 |  |
| Gorzs4 | Tisza | Hódmezővásárhely-Gorzsa | 46.4 | 20.4 | 5000–4500 | F | T1a | .. | 0.06 | 11.4±3.1 | 23±11 |  |
| KOKE3a | Tisza | Hódmezővásárhely-Kökénydomb Vörös tanya | 46.4 | 20.2 | 5000–4500 | M | K1b1 | I | 0.06 | 13.9±3.2 | ± |  |
| PULE1.24 | Tisza | Pusztataskony-Ledence | 47.5 | 20.5 | 5000–4500 | F | K1a4 | .. | 0.40 | 10.5±1.9 | 18±7.1 |  |
| VSM3a | Tisza | Vesztő-Mágor | 46.9 | 21.2 | 5000–4500 | M | H26 | G2a | 0.09 | 7.00±2.6 | ± |  |
| ALE14a | TDLN | Alsónyék-Elkerülő site 2 | 46.2 | 18.7 | 5030–4848 | M | U8b1b | G2a | 0.05 | -1.12±3.2 | ± |  |
| ALE4a | TDLN | Alsónyék-Elkerülő site 2 | 46.2 | 18.7 | 5016–4838 | M | T2c1 | F | 0.03 | 10.7±3.6 | ± |  |
| BAL3a | TDLN | Bátaszék-Lajvér | 46.2 | 18.7 | 4800–4500 | M | T2f | H1b1 | 0.91 | 6.97±1.7 | 22±9.1 |  |
| CSAT19a | TDLN | Csabdi-Télizöldes | 47.5 | 18.6 | 4800–4500 | M | H | H | 0.52 | 5.89±1.8 | 34±9.6 |  |
| CSAT25a | TDLN | Csabdi-Télizöldes | 47.5 | 18.6 | 4826–4602 | M | T2b | I2 | 0.43 | 13.7±1.9 | 26±8.1 |  |
| FAGA1a | TDLN | Fajsz-Garadomb | 46.4 | 18.9 | 5100–4750 | M | HV0a | I | 0.09 | 5.13±2.4 | ± |  |
| FAGA2a | TDLN | Fajsz-Garadomb | 46.4 | 18.9 | 5195–4842 | F | H | .. | 0.49 | 12.1±1.8 | 14±4.1 |  |
| FEB3a | TDLN | Felsőörs-Bárókert | 47.0 | 18.0 | 4800–4500 | M | H44 | J2a | 0.16 | 6.38±2.1 | ± |  |
| HUNG347, NE7 | TDLN | Apc-Berekalja | 47.2 | 19.8 | 4491–4357 | M | N1a1a1a | I | 4.85 | 10.7±1.6 | 19±3.1 | [12, 22] |
| SZEH5a | TDLN | Szemely-Hegyes | 46.0 | 18.3 | 4904–4709 | M | K1b1a | G | 0.01 | 10.9±6.6 | ± |  |
| SZEH7b | TDLN | Szemely-Hegyes | 46.0 | 18.3 | 4930–4715 | F | K1a | .. | 0.52 | 3.48±1.7 | ± |  |
| VEJ12a | TDLN | Veszprém Jutasi út | 47.1 | 17.9 | 4800–4500 | M | U8b1a2b | H | 0.10 | 6.24±2.3 | ± |  |
| VEJ2a | TDLN | Veszprém Jutasi út | 47.1 | 17.9 | 4800–4500 | M | T2b | C | 0.34 | 5.69±1.8 | ± |  |
| VEJ5a | TDLN | Veszprém Jutasi út | 47.1 | 17.9 | 4800–4500 | M | J1c2 | G2a2a1 | 0.62 | 7.87±1.8 | 15±2.9 |  |
| GEN67 | Tiszapolgár | Törökszentmiklós road 4 site 3 | 47.2 | 20.4 | 4444–4257 | M | H1 | I2a2a1b | 2.28 | 13.1±1.7 | 50±15 |  |
| PULE1.10a | Tiszapolgár | Pusztataskony-Ledence | 47.5 | 20.5 | 4500–4000 | M | T2c1 | I2a | 0.28 | 9.14±2.0 | ± |  |
| PULE1.13a | Tiszapolgár | Pusztataskony-Ledence | 47.5 | 20.5 | 4500–4000 | M | T2c1 | G2a2b2a1a1c1a | 0.38 | 10.4±1.9 | ± |  |
| PULE1.9a | Tiszapolgár | Pusztataskony-Ledence | 47.5 | 20.5 | 4500–4000 | M | H26 | G2a2b | 0.11 | 11.7±2.4 | ± |  |
| GEN100 | Lásinja | Alsónyék, site 11 | 46.2 | 18.7 | 4300–3900 | F | T2b | .. | 1.81 | 9.62±1.6 | 45±11 |  |
| GEN49 | Lásinja | Nemesnádudvar-Papföld | 46.3 | 19.1 | 4228–3963 | M | T2b23 | CT | 0.97 | 12.9±1.8 | 27±6.8 |  |
| KEFP2a | Lásinja | Keszthely-Fenékpuszta | 46.7 | 17.2 | 4300–3900 | F | J2b1a | .. | 0.74 | 9.23±1.7 | 21±5.4 |  |
| KON2a | Lásinja | Enese elkerülő, Kóny, Proletár-dűlő, M85, Site 2 | 47.6 | 17.4 | 4333–4072 | F | K2a | .. | 2.13 | 10.4±1.7 | 21±6.4 |  |
| M6-116.12a | Lásinja | Lánycsók, Csata-alja | 46.0 | 18.6 | 4232–4046 | F | T2f8a | .. | 0.64 | 9.79±1.7 | 29±11 |  |
| VEJ9a | Lásinja | Veszprém Jutasi út | 47.1 | 17.9 | 4339–4237 | M | H40 | CT | 0.05 | 8.93±3.2 | ± |  |
| GEN60 | Protoboleráz | Abony, Turjános-dűlő | 47.2 | 20.0 | 3909–3651 | M | H | G2a2b2a | 1.88 | 14.2±1.6 | 37±8.8 |  |
| GEN61 | Protoboleráz | Abony, Turjános-dűlő | 47.2 | 20.0 | 3800–3600 | M | J1c | I2c | 0.76 | 10.9±1.7 | 65±14 |  |
| GEN62 | Protoboleráz | Abony, Turjános-dűlő | 47.2 | 20.0 | 3800–3600 | F | N1a1a1a3 | .. | 4.81 | 8.10±1.6 | 37±9.6 |  |
| GEN63 | Protoboleráz | Abony, Turjános-dűlő | 47.2 | 20.0 | 3762–3636 | M | U5a1c1 | I2c | 1.92 | 12.0±1.7 | 34±8.1 |  |
| GEN12a | Baden | Budakalász-Luppa csárda | 47.6 | 19.0 | 3300–2850 | M | H26a | G2a2b2a1a1b1 | 1.98 | 14.0±1.7 | 34±7.2 |  |
| GEN13a | Baden | Budakalász-Luppa csárda | 47.6 | 19.0 | 3300–2850 | M | HV | G2a2b2a1a | 2.65 | 11.4±1.6 | 27±6.6 |  |
| GEN15a | Baden | Budakalász-Luppa csárda | 47.6 | 19.0 | 3367–3103 | M | J2a1a1 | G2a2b2a1a1c1a | 1.66 | 10.9±1.7 | 22±9.2 |  |
| GEN16a | Baden | Alsónémedi | 47.3 | 19.2 | 3300–2850 | F | T2b | .. | 4.30 | 13.0±1.6 | 38±16 |  |
| GEN17a | Baden | Alsónémedi | 47.3 | 19.2 | 3359–3098 | M | U5b3f | G2a2a | 0.82 | 10.9±1.7 | 21±6.3 |  |
| GEN21 | Baden | Balatonlelle-Felső-Gamász | 46.8 | 17.7 | 3600–2850 | M | K1a | I2a1 | 0.67 | 12.4±1.7 | ± |  |
| GEN22 | Baden | Balatonlelle-Felső-Gamász | 46.8 | 17.7 | 3600–2850 | M | U5a1 | I2a1a1 | 2.31 | 14.7±1.7 | 25±6.6 |  |
| GEN55 | Baden | Vámosgyörk | 47.7 | 19.9 | 3600–2850 | F | T2c2 | .. | 0.81 | 13.2±1.8 | 22±6.7 |  |
| HUNG353, CO1 | Baden | Apc-Berekalja | 47.2 | 19.8 | 3315–2923 | F | H | .. | 4.56 | 15.3±1.7 | 37±15 | [12, 22] |
| Vors1 | Baden | Vörs | 46.7 | 17.3 | 3300–2850 | F | T2f | .. | 0.03 | 4.52±4.2 | ± |  |

Extended Data Table 1 – continued from previous page
| ID | Population | Site | Lat. | Long. | Date | Sex | Mt Hap | Y Hap | Cov. | HG% | ALD | Ref. |
| --- | --- | --- | --- | --- | --- | --- | --- | --- | --- | --- | --- | --- |
| HAL03a | LBK | Halberstadt-Sonntagsfeld | 51.9 | 11.0 | 5295–5057 | F | T2b | .. | 0.01 | -5.19±6.8 | ± |  |
| HAL07a | LBK | Halberstadt-Sonntagsfeld | 51.9 | 11.0 | 5212–4992 | F | N1a1b | .. | 0.05 | 1.74±3.2 | ± |  |
| HAL15a | LBK | Halberstadt-Sonntagsfeld | 51.9 | 11.0 | 5199–4857 | M | N1a1a3 | G2 | 0.02 | 5.32±5.0 | ± |  |
| HAL17b | LBK | Halberstadt-Sonntagsfeld | 51.9 | 11.0 | 5500–4850 | F | V1 | .. | 0.02 | 9.32±4.3 | ± |  |
| HAL18a | LBK | Halberstadt-Sonntagsfeld | 51.9 | 11.0 | 5500–4850 | F | K2a | .. | 0.02 | 0.27±4.7 | ± |  |
| HAL19 | LBK | Halberstadt-Sonntagsfeld | 51.9 | 11.0 | 5500–4850 | F | K1a2 | .. | 0.86 | 7.18±1.7 | 17±5.6 | [12]* |
| HAL2 | LBK | Halberstadt-Sonntagsfeld | 51.9 | 11.0 | 5211–4963 | M | N1a1a1a2 | G2a2a1 | 0.76 | 1.93±1.7 | 10±2.3 | [10, 12]* |
| HAL20b | LBK | Halberstadt-Sonntagsfeld | 51.9 | 11.0 | 5500–4850 | M | K1a2 | G2a2a | 0.06 | 2.56±3.1 | ± |  |
| HAL21a | LBK | Halberstadt-Sonntagsfeld | 51.9 | 11.0 | 5500–4850 | M | T2b | G2a2a | 0.01 | -4.46±5.9 | ± |  |
| HAL22b | LBK | Halberstadt-Sonntagsfeld | 51.9 | 11.0 | 5500–4850 | F | T2b | .. | 0.02 | -7.80±4.8 | ± |  |
| HAL24 | LBK | Halberstadt-Sonntagsfeld | 51.9 | 11.0 | 5201–4850 | M | X2d1 | G2a2a1 | 0.42 | 6.47±1.8 | ± | [10, 12]* |
| HAL25 | LBK | Halberstadt-Sonntagsfeld | 51.9 | 11.0 | 5210–5002 | M | K1a | G2a2a1 | 0.49 | 2.61±1.7 | 17±7.5 | [10, 12]* |
| HAL27a | LBK | Halberstadt-Sonntagsfeld | 51.9 | 11.0 | 5500–4850 | M | N1a1a3 | G2a2a | 0.05 | 3.88±3.0 | ± |  |
| HAL31a | LBK | Halberstadt-Sonntagsfeld | 51.9 | 11.0 | 5295–5057 | F | K1c | .. | 0.12 | 4.59±2.3 | 10±3.1 |  |
| HAL32b | LBK | Halberstadt-Sonntagsfeld | 51.9 | 11.0 | 5500–4850 | F | H26 | .. | 0.23 | 3.37±2.0 | ± |  |
| HAL34 | LBK | Halberstadt-Sonntagsfeld | 51.9 | 11.0 | 5219–5021 | F | N1a1a1 | .. | 0.25 | 5.69±2.0 | 10±4.0 | [10, 12]* |
| HAL35b | LBK | Halberstadt-Sonntagsfeld | 51.9 | 11.0 | 5500–4850 | F | J1c | .. | 0.10 | 3.98±2.4 | ± |  |
| HAL38a | LBK | Halberstadt-Sonntagsfeld | 51.9 | 11.0 | 5500–4850 | F | V1 | .. | 0.29 | 1.12±1.9 | ± |  |
| HAL39b | LBK | Halberstadt-Sonntagsfeld | 51.9 | 11.0 | 5210–5002 | M | H1e | G2a2a1 | 0.08 | 4.01±2.7 | ± |  |
| HAL4 | LBK | Halberstadt-Sonntagsfeld | 51.9 | 11.0 | 5202–4852 | F | N1a1a1a | .. | 6.92 | 6.62±1.6 | 18±5.8 | [10, 12]* |
| HAL40a | LBK | Halberstadt-Sonntagsfeld | 51.9 | 11.0 | 5500–4850 | F | T2b | .. | 0.17 | 2.53±2.1 | ± |  |
| HAL5 | LBK | Halberstadt-Sonntagsfeld | 51.9 | 11.0 | 5211–4991 | F | T2c1 | .. | 2.23 | 3.01±1.6 | 10±6.9 | [10, 12]* |
| KAR16A | LBK | Karsdorf | 51.3 | 11.7 | 5500–4850 | M | H46b | T1a | 0.09 | 0.29±2.7 | 13±5.1 | [12] |
| KAR6 | LBK | Karsdorf | 51.3 | 11.7 | 5217–5041 | M | H1/H1a1b | CT | 0.10 | 5.85±2.6 | ± | [10, 12] |
| LBK1976 | LBK | Viesenhäuser Hof | 48.8 | 9.2 | 5500–4850 | F | T2e | .. | 0.44 | 3.50±1.7 | 18±4.4 | [10, 12] |
| LBK1992 | LBK | Viesenhäuser Hof | 48.8 | 9.2 | 5500–4850 | F | T2b | .. | 2.66 | 5.75±1.6 | 12±4.3 | [10, 12] |
| LBK2155 | LBK | Viesenhäuser Hof | 48.8 | 9.2 | 5500–4850 | F | T2b | .. | 3.63 | 4.89±1.5 | 13±4.4 | [10, 12] |
| Stuttgart | LBK | Viesenhäuser Hof | 48.8 | 9.2 | 5310–5076 | F | T2c1b | .. | 9.65 | 3.04±1.6 | 22±8.1 | [9]* |
| UWS4 | LBK | Unterwiederstedt | 51.7 | 11.5 | 5223–5021 | F | J1c17 | .. | 18.6 | 5.76±1.6 | 13±14 | [10, 12] |
| ESP30 | GermanyMN | Esperstedt | 51.4 | 11.7 | 3970–3710 | M | H1e1a | I | 0.09 | 22.2±2.7 | ± | [10, 12] |
| HAL13a | GermanyMN | Halberstadt-Sonntagsfeld | 51.9 | 11.0 | 4600–4300 | F | V1a | .. | 0.11 | 9.14±2.4 | 13±4.3 |  |
| QLB15D | GermanyMN | Quedlinburg | 51.8 | 11.1 | 3654–3527 | M | HV | R | 0.16 | 21.1±2.2 | 36±8.7 | [10, 12] |
| QLB18A | GermanyMN | Quedlinburg | 51.8 | 11.1 | 3640–3376 | F | T2e1 | .. | 0.41 | 19.8±1.8 | 23±4.9 | [10, 12] |
| SALZ3B | GermanyMN | Salzmünde-Schiebzig | 51.5 | 11.8 | 3400–3025 | M | U3a1 | G2a2a1 | 0.09 | 15.1±2.7 | ± | [12] |
| SALZ57A | GermanyMN | Salzmünde-Schiebzig | 51.5 | 11.8 | 3345–3097 | F | H3 | .. | 0.02 | 25.3±4.5 | ± |  |
| SALZ77A | GermanyMN | Salzmünde-Schiebzig | 51.5 | 11.8 | 3400–3025 | M | H3 | IJK (x J) | 0.02 | 21.6±5.1 | ± |  |
| Bla16 | Blätterhöhle | Blätterhöhle Cave | 51.4 | 7.6 | 3958–3344 | M | U5b2a2 | R1b1 | 0.80 | 39.9±2.0 | 15±5.8 |  |
| Bla28 | Blätterhöhle | Blätterhöhle Cave | 51.4 | 7.6 | 3337–3024 | M | J1c1b1 | R1 | 0.10 | 52.5±2.7 | 11±4.5 |  |
| Bla5 | Blätterhöhle | Blätterhöhle Cave | 51.4 | 7.6 | 3704–3117 | F | H5 | .. | 5.07 | 41.6±1.9 | 24±4.7 |  |
| Bla8 | Blätterhöhle | Blätterhöhle Cave | 51.4 | 7.6 | 4038–3532 | M | U5b2b2 | I2a1 | 4.58 | 73.4±2.0 | 12±2.9 |  |
| CB13 | Iberia EN | Cova Bonica | 41.4 | 1.9 | 5469–5327 | F | K1a2a | .. | 0.98 | 10.1±1.8 | 17±3.5 | [23] |
| E-06-Ind1 | Iberia EN | El Prado de Pancorbo | 42.6 | -3.1 | 4827–4692 | F | K1a4a1 | .. | 0.47 | 8.82±1.9 | 17±2.3 |  |
| E-14-Ind2 | Iberia EN | El Prado de Pancorbo | 42.6 | -3.1 | 5216–5031 | F | H1 | .. | 0.38 | 7.60±1.8 | 19±2.8 |  |
| Troc1 | Iberia EN | Els Trocs | 42.5 | 0.5 | 5311–5218 | F | J1c3 | .. | 0.69 | 7.23±1.8 | 12±9.0 | [10, 12] |
| Troc3 | Iberia EN | Els Trocs | 42.5 | 0.5 | 5294–5066 | M | T2c1d/T2c1d2 | R1b1a2 | 1.31 | 10.0±1.8 | 49±22 | [10, 12] |
| Troc5 | Iberia EN | Els Trocs | 42.5 | 0.5 | 5310–5078 | M | N1a1a1 | I2a1b1 | 13.8 | 6.91±1.6 | 6.8±2.8 | [10, 12] |
| Troc7 | Iberia EN | Els Trocs | 42.5 | 0.5 | 5303–5075 | F | V | .. | 1.57 | 11.2±1.7 | 18±4.8 | [10, 12] |
| Mina18 | Iberia MN | La Mina | 41.3 | -2.3 | 3893–3661 | F | U5b1 | .. | 13.6 | 23.1±1.8 | 42±18 | [10, 12] |
| Mina3 | Iberia MN | La Mina | 41.3 | -2.3 | 3900–3600 | M | K1a1b1 | H2 | 0.38 | 19.7±1.9 | 80±20 | [10, 12] |
| Mina4 | Iberia MN | La Mina | 41.3 | -2.3 | 3900–3600 | M | H1 | I2a2a1b2 | 3.95 | 22.8±1.9 | 25±6.3 | [10, 12] |
| Mina6 | Iberia MN | La Mina | 41.3 | -2.3 | 3900–3600 | F | K1b1a1 | .. | 1.36 | 19.1±1.7 | 46±8.1 | [10, 12] |
| 1.-K11 | Iberia CA | La Chabola de la Hechicera | 42.6 | -2.6 | 3263–2903 | M | X2b | I2a2 | 0.18 | 28.1±2.1 | 67±25 |  |
| 3.-K11 | Iberia CA | La Chabola de la Hechicera | 42.6 | -2.6 | 3627–3363 | F | J2a1a1 | .. | 0.12 | 24.7±2.5 | 30±12 |  |
| 5.-K18 | Iberia CA | La Chabola de la Hechicera | 42.6 | -2.6 | 3090–2894 | M | J1c1 | I2a2 | 0.10 | 18.7±2.5 | 43±11 |  |
| ES.1/4 | Iberia CA | El Sotillo | 42.6 | -2.6 | 2571–2347 | M | H3 | I | 0.07 | 25.6±2.8 | ± |  |
| ES-6G-110 | Iberia CA | El Sotillo | 42.6 | -2.6 | 2916–2714 | M | H3 | I2a2a | 0.05 | 25.7±3.2 | ± |  |
| Inventario0/4 | Iberia CA | El Sotillo | 42.6 | -2.6 | 2481–2212 | M | X2b | I2a2a | 0.12 | 30.0±2.5 | 56±23 |  |
| LHUE11J.5 | Iberia CA | Alto de la Huesera | 42.6 | -2.6 | 3092–2877 | F | U5b1 | .. | 1.19 | 27.0±1.9 | 40±9.7 |  |
| LHUE2010.10 | Iberia CA | Alto de la Huesera | 42.6 | -2.6 | 3014–2891 | F | J1c1 | .. | 0.11 | 25.5±2.5 | 64±13 |  |
| LHUE2010.11 | Iberia CA | Alto de la Huesera | 42.6 | -2.6 | 3092–2918 | M | V | G2a2a | 0.32 | 31.5±2.0 | ± |  |
| LHUE2014.11J | Iberia CA | Alto de la Huesera | 42.6 | -2.6 | 3100–2850 | F | U5b2b | .. | 0.06 | 26.6±3.0 | ± |  |
| LY.II.A.10.15066 | Iberia CA | Las Yurdinas II | 42.6 | -2.7 | 3350–2750 | M | U5b2b3a | I2a2a | 1.93 | 30.4±1.8 | ± |  |
| LY.II.A.10.15067 | Iberia CA | Las Yurdinas II | 42.6 | -2.7 | 3350–2750 | F | J2a1a1 | .. | 0.30 | 24.1±2.0 | ± |  |
| LY.II.A.10.15068 | Iberia CA | Las Yurdinas II | 42.6 | -2.7 | 3350–2750 | F | K1a4a1a | .. | 0.39 | 29.5±2.0 | 28±10 |  |
| LY.II.A.10.15069 | Iberia CA | Las Yurdinas II | 42.6 | -2.7 | 3354–2943 | F | J1c3 | .. | 4.24 | 25.4±1.7 | 29±15 |  |
| MIR1 | Iberia CA | El Mirador Cave | 42.3 | -3.5 | 2900–2346 | F | K1a | .. | 0.24 | 24.5±2.1 | ± | [12] |
| MIR13 | Iberia CA | El Mirador Cave | 42.3 | -3.5 | 2900–2346 | F | H3c3 | .. | 0.10 | 28.1±2.4 | ± | [12] |
| MIR14 | Iberia CA | El Mirador Cave | 42.3 | -3.5 | 2568–2346 | M | H3 | I2a2a | 0.94 | 23.5±1.8 | 57±16 | [12] |
| MIR17 | Iberia CA | El Mirador Cave | 42.3 | -3.5 | 2900–2346 | F | J1c1 | .. | 0.22 | 23.8±2.2 | ± | [12] |
| MIR18 | Iberia CA | El Mirador Cave | 42.3 | -3.5 | 2865–2575 | F | H1t | .. | 1.58 | 20.2±1.6 | 43±17 | [12] |
| MIR19 | Iberia CA | El Mirador Cave | 42.3 | -3.5 | 2900–2346 | M | H3 | I | 0.06 | 22.1±3.1 | ± | [12] |
| MIR2 | Iberia CA | El Mirador Cave | 42.3 | -3.5 | 2857–2496 | F | K1b1a1 | .. | 0.98 | 22.9±1.8 | 56±9.0 | [12] |
| MIR21 | Iberia CA | El Mirador Cave | 42.3 | -3.5 | 2900–2346 | M | H3 | I | 0.11 | 25.0±2.4 | 55±17 | [12] |
| MIR22 | Iberia CA | El Mirador Cave | 42.3 | -3.5 | 2900–2346 | F | K1a2a | .. | 2.79 | 22.9±1.7 | 62±10 | [12] |
| MIR24 | Iberia CA | El Mirador Cave | 42.3 | -3.5 | 2900–2346 | M | J2b1a3 | G2a2b2b | 0.06 | 20.3±3 | ± | [12] |
| MIR25 | Iberia CA | El Mirador Cave | 42.3 | -3.5 | 2900–2346 | M | U3a1 | I2a1b1 | 0.73 | 25.5±1.7 | ± | [12] |
| MIR5, MIR6 | Iberia CA | El Mirador Cave | 42.3 | -3.5 | 2900–2679 | M | X2b | C1a2 | 10.4 | 21.0±1.7 | 107±41 | [12] |

Cov: average coverage per SNP. HG%: inferred percentage of hunter-gatherer ancestry (mean ± standard error). ALD: inferred date of admixture (generations in the past; mean ± standard error). Ref: reference for published data; if blank, newly published sample in this study (asterisk denotes a published individual with new sequencing data added). Radiocarbon dates are in normal text, while dates estimated from archaeological context are in italics. Further information can be found in Supplementary Table 1.

A principal component analysis (PCA [26]) of our samples showed that, as expected, all of the Neolithic individuals fall along a cline of admixture between FEF and WHG, with varying proportions of hunter-gatherer ancestry (Extended Data Figure 1). All studied population groups are consistent with a common origin in Anatolia (Supplementary Information section 4), and differentiation among the ancestral farmer populations in the three regions is low (Extended Data Figure 1). To investigate genetic structure within the source population(s) of hunter-gatherer ancestry, we combined the four primary WHG individuals with other hunter-gatherers (eastern hunter-gatherers, or “EHG,” ∼7000–5000 BCE from Russia [10,12]; “Bichon,” ∼10,000 BCE from Switzerland [27]; and “El Mirón,” ∼17,000 BCE from Spain [25]) along an approximate east–west transect across Europe. We computed pairwise outgroup *f*_3_-statistics, performed multidimensional scaling (MDS) on the resulting matrix, and converted the MDS positions to polar coordinates (Supplementary Information section 5). The correlation between genetic structure and longitude revealed by this analysis (Figure 1D) suggests that our reference samples can reasonably be used to define a geographic cline among hunter-gatherers within Europe. Computing *f*_4_-statistics measuring shared drift between our Neolithic samples and the WHG individuals, we observed an increasing trend in hunter-gatherer ancestry over time in each region [10–12], although at a slower rate in Hungary than in Germany and Spain, and with limited intra-population structure or heterogeneity (Figure 2A; Supplementary Information section 6). We find that this hunter-gatherer ancestry is more similar to the eastern WHG individuals (KO1 and Villabruna) farther east and more similar to the western WHG individuals (LB1 and Loschbour) farther west (Figure 2B). While this pattern does not demonstrate directly where mixture between hunter-gatherers and farmers took place, it suggests that hunter-gatherer ancestry in farmers was to a substantial extent derived from populations from relatively close to where they lived.

**Figure 2.**
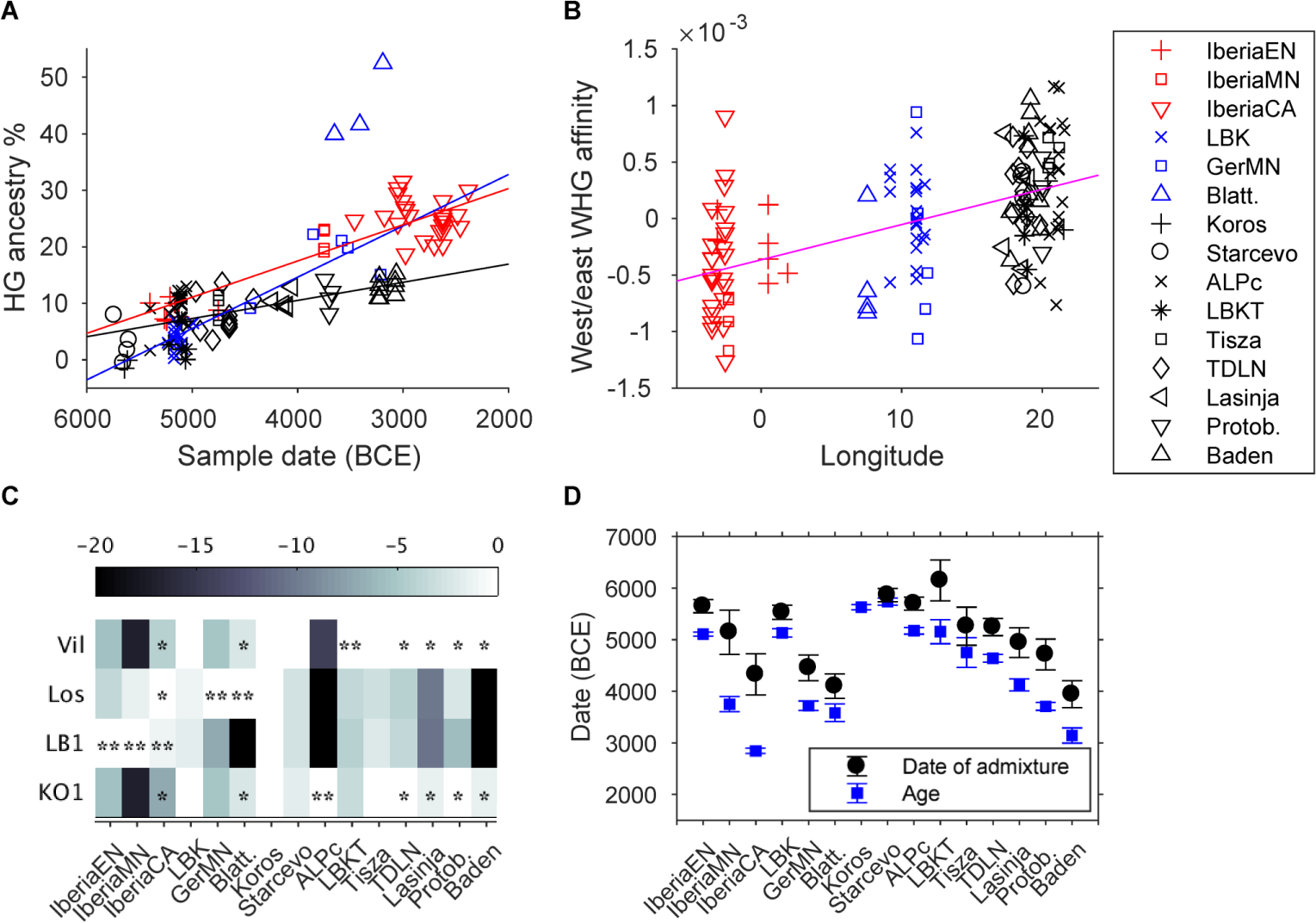
Admixture parameters for test individuals and populations. (A) Estimated individual hunter-gatherer ancestry versus sample date, with best-fitting regression lines for each region (excluding Blätterhöhle). Standard errors are around 2% for hunter-gatherer ancestry and 100 years for dates (Methods; Extended Data Table 1). Individual Bla8 (∼74% hunter-gatherer ancestry) is omitted for scale. (B) Relative affinity of hunter-gatherer ancestry in Neolithic individuals, measured as *f*_4_(LB1+Loschbour, KO1+Villabruna; Anatolia, *X*) (positive, more similar to eastern WHG; negative, more similar to western WHG; standard errors ∼5×10^−4^), with best-fitting regression line (nominal *p* ∼ 10^−11^, unadjusted for non-independence among samples). Within-region correlations are not significant (Hungary *p* ∼ 0.1), whereas the three regions in aggregate differ from each other at |*Z*| > 3. (C) Relative log-likelihood of admixture graph models fitting each population as a mixture of FEF and one of the four WHG individuals (Los, Loschbour; Vil, Villabruna). Asterisks denote statistical significance (*p* < 0.05; Supplementary Information section 6), with single asterisks for significant components whose source is not uniquely identified (Extended Data Table 2). (D) Population-level average sample ages and dates of admixture, plus or minus two standard errors. Due to heterogeneity, we omit the outlier individuals Bla28 (Blätterhöhle) and GEN61 (Protoboleráz) in the dates.

**Extended Data Figure 1.**
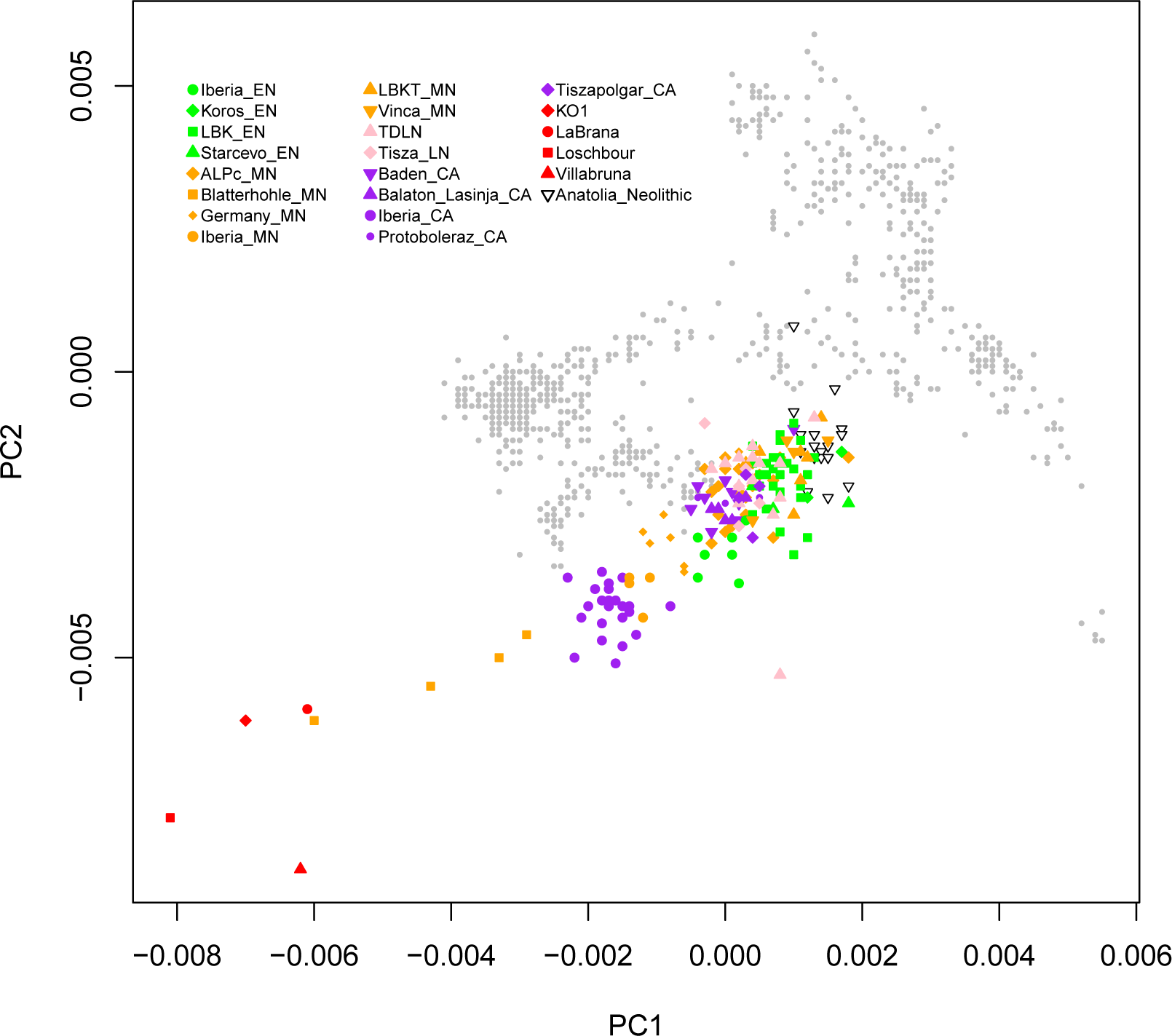
First two principal components from PCA. We merged with populations genotyped on the Affymetrix Human Origins array and computed PCs for a set of 826 present-day western Eurasian individuals (background gray points). We then projected ancient individuals using the “lsqproject” and “shrinkmode” options in smartpca [26]. Shown is a closeup omitting the present-day Bedouin individuals.

To analyze admixed hunter-gatherer ancestry more formally, we modeled Neolithic farmers in an admixture graph framework. We started with a “scaffold” model (Extended Data Figure 3) consisting of Neolithic Anatolians, the four reference WHG individuals, and two outgroups (Mbuti and Kostenki 14 [25, 28]). We observed discrete signals of admixture in LB1 and KO1 via *f*_3_-and *f*_4_-statistics [29], and both fit best as admixed in the scaffold model, LB1 with ancestry from a deeper European hunter-gatherer lineage and KO1 with a small proportion of FEF admixture (Supplementary Information section 6). We then added each Neolithic population to this model in turn, fitting them as a mixture of FEF and either one or two hunter-gatherer ancestry components. To check for robustness, we repeated our analyses using transversions or outgroup-ascertained SNPs only, with in-solution capture data for Loschbour, and with additional or alternative hunter-gatherers in the model (Extended Data Table 2; Supplementary Information section 6), and in all cases the results were qualitatively consistent. We find that almost all ancient groups from Hungary have ancestry significantly closest to one of the more eastern WHG individuals (either KO1 or Villabruna); the samples from present-day Germany have greatest affinity to Loschbour; and all three Iberian groups contain LB1-related ancestry (Figure 2C; Extended Data Table 2). This pattern implies that admixture into European farmers occurred multiple times from local hunter-gatherer populations. Moreover, combining the proportions and sources of hunter-gatherer ancestry, populations from the three regions are distinguishable at all stages of the Neolithic. Thus, any further migrations that may have occurred after the initial spread of farming were not substantial enough within the studied regions to disrupt the observed heterogeneity.

**Extended Data Figure 3.**
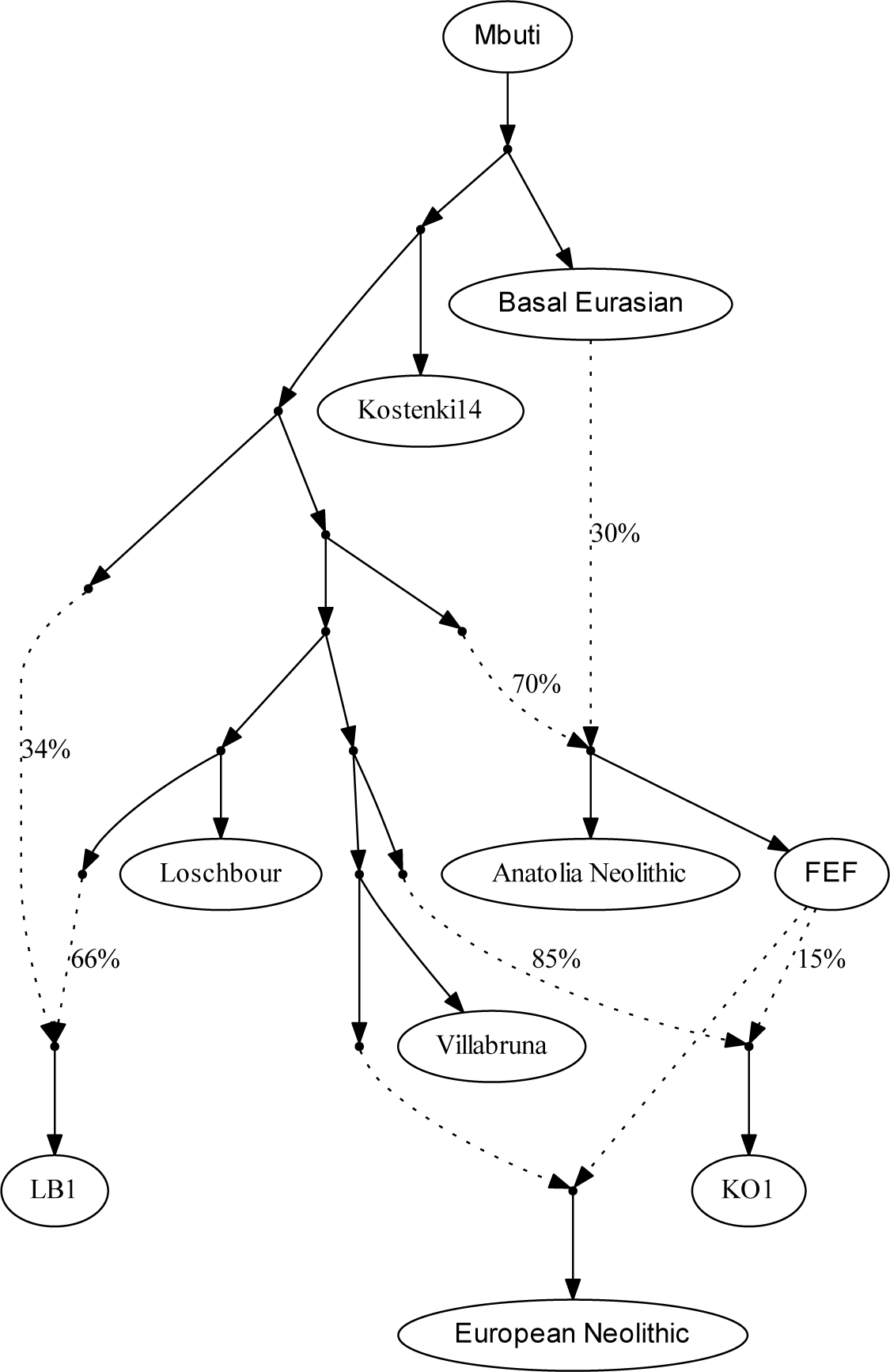
Scaffold admixture graph used for modeling European Neolithic populations. Dotted lines denote admixture events. Anatolia Neolithic, LB1, and KO1 are modeled as admixed, with Basal Eurasian ancestry, deeper European hunter-gatherer ancestry, and FEF ancestry, respectively. European test populations are fit as a mixture of FEF and ancestry related to one or two of the four WHG individuals (here Villabruna-related as an example). See Supplementary Information section 6 for full details.

**Extended Data Table 2.**
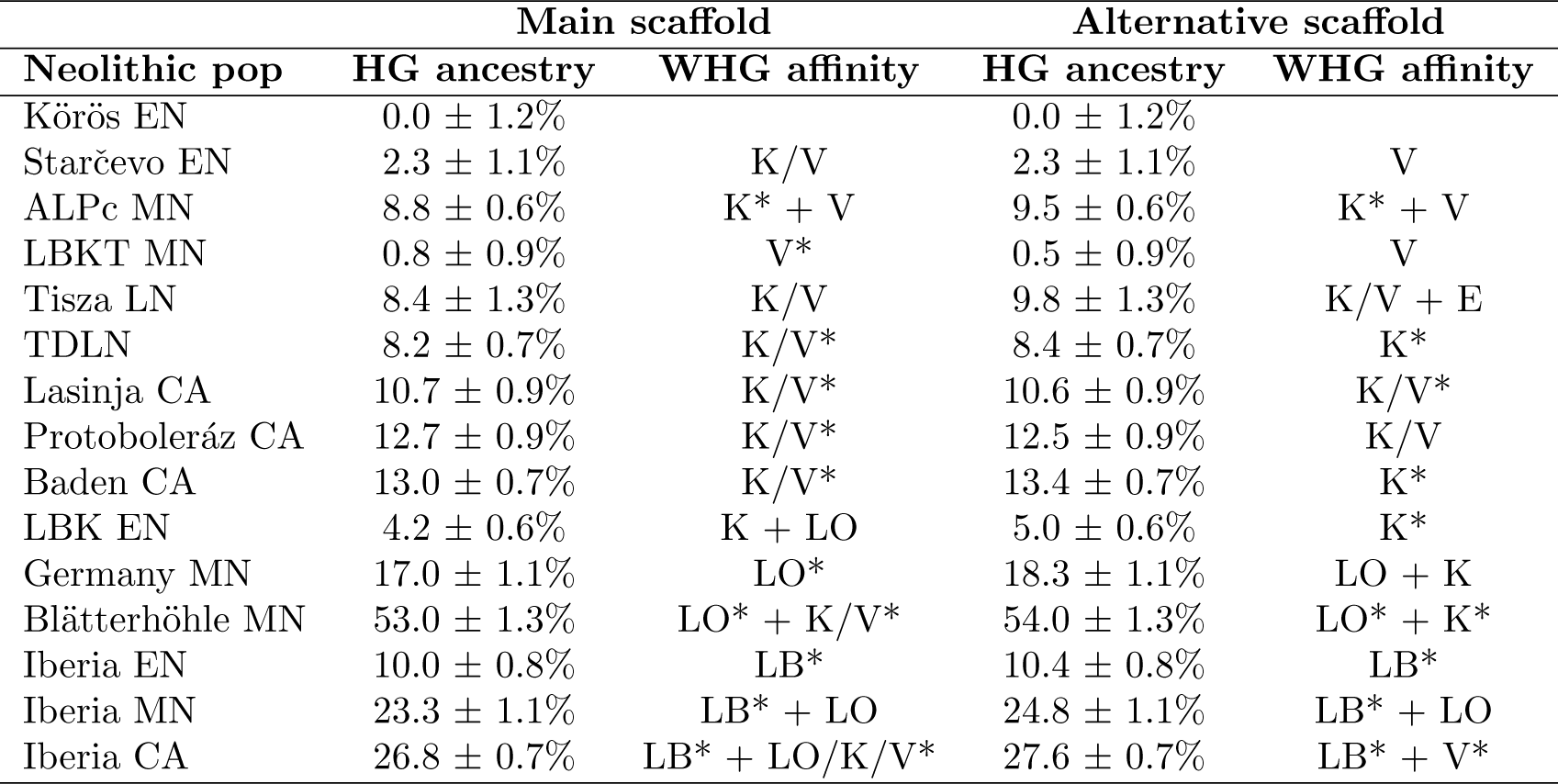
Admixture graph results for Neolithic populations

Additional insights about population interactions can be gained by studying the dates of admixture events. We used *ALDER* [30] to estimate dates of admixture for Neolithic individuals based on the recombination-induced breakdown of contiguous blocks of FEF and WHG ancestry over time (Extended Data Table 1). The *ALDER* algorithm is not able to accommodate large amounts of missing data, so we developed a strategy for running it with the relatively low coverage of ancient DNA (Supplementary Information section 7). To obtain calendar dates of admixture (Figure 2D), we combine the *ALDER* results (in generations in the past) with the ages of the Neolithic individuals, assuming an average generation time of 28 years [31,32]. These dates are based on a model of a single wave of admixture, which means that if the true history for a population includes multiples waves or continuous admixture, we will obtain an intermediate value. Additionally, while the primary signal is due to admixture between farmers and hunter-gatherers, mixture among farmers can also be detected if the groups have different proportions of hunter-gatherer ancestry, meaning that inferred dates, especially for later populations, may not all reflect admixture between farmers and unadmixed hunter-gatherers.

For our most complete time series, from Hungary, we infer admixture dates throughout the Neolithic that are on average mostly 18–30 generations old (500–840 years), indicating a degree of ongoing population transformation and admixture ((Figure 2D; Extended Data Table 3). This pattern is accompanied by a gradual increase in hunter-gatherer ancestry over time, although never reaching the levels observed in MN Germany or Iberia (Figure 2A). While five of the six EN individuals from Hungary do not have significantly more hunter-gatherer ancestry than Neolithic Anatolians (Figure 2A; Extended Data Table 1), one Starčevo sample, BAM17b, is inferred to have 7.9 ± 1.7% hunter-gatherer ancestry and a very recent *ALDER* date of 4.5 ± 1.9 generations (5865 ± 65 BCE; 2.9 ± 1.6 generations using a group-level estimate; Extended Data Table 3), consistent with his having had one or two hunter-gatherer ancestors in the last several generations. We also infer an average admixture date of 5700 ± 65 BCE for the ALPc MN, again suggesting that in Hungary, interaction between Anatolian migrants and local HGs began in the Early Neolithic (cf. [14,33–35]). The greatest differences between Alföld and Transdanubia are observed in the MN, with substantially more hunter-gatherer ancestry in ALPc than LBKT (Figure 2; Extended Data Table 2), and overall, we observe slight trends toward more hunter-gatherer ancestry to the north and east (Extended Data Figure 2), as expected based on the greater archaeological evidence of hunter-gatherer settlement and interactions [33]. By the LN and CA, however, and especially in the Baden period (when the region became culturally unified [36]) our results are broadly similar over the two halves of present-day Hungary.

**Extended Data Figure 2.**
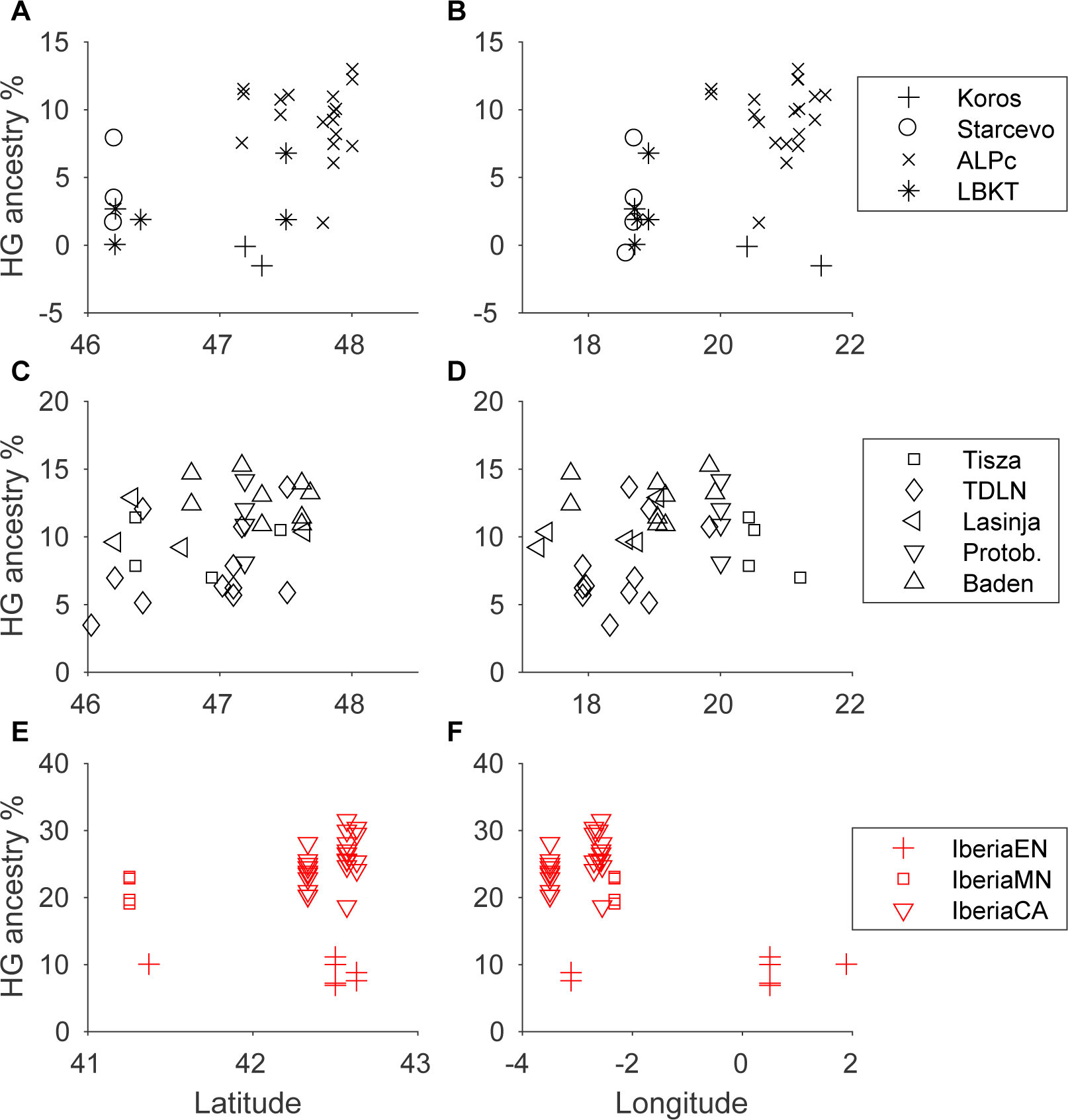
Hunter-gatherer ancestry as a function of latitude and longitude for Neolithic individuals from (A), (B) EN/MN Hungary; (C), (D) LN/CA Hungary; and (E), (F) Iberia. Protob., Protoboleráz.

**Extended Data Table 3.**
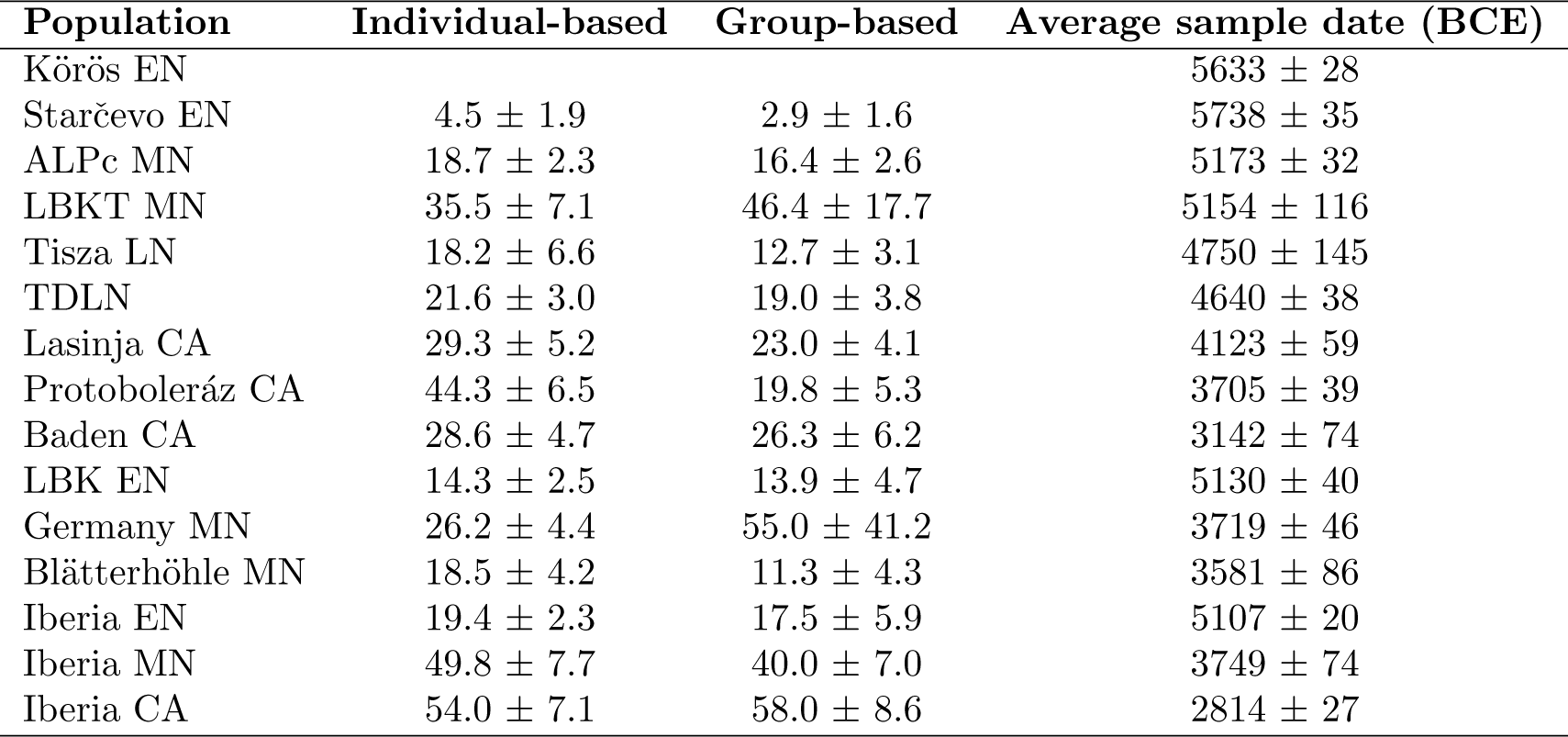
Average dates of admixture for Neolithic populations

| Population | Individual-based | Group-based | Average sample date (BCE) |
| --- | --- | --- | --- |
| Körös EN | | | 5633 ± 28 |
| Starčevo EN | 4.5 ± 1.9 | 2.9 ± 1.6 | 5738 ± 35 |
| ALPc MN | 18.7 ± 2.3 | 16.4 ± 2.6 | 5173 ± 32 |
| LBKT MN | 35.5 ± 7.1 | 46.4 ± 17.7 | 5154 ± 116 |
| Tisza LN | 18.2 ± 6.6 | 12.7 ± 3.1 | 4750 ± 145 |
| TDLN | 21.6 ± 3.0 | 19.0 ± 3.8 | 4640 ± 38 |
| Lasinja CA | 29.3 ± 5.2 | 23.0 ± 4.1 | 4123 ± 59 |
| Protoboleráz CA | 44.3 ± 6.5 | 19.8 ± 5.3 | 3705 ± 39 |
| Baden CA | 28.6 ± 4.7 | 26.3 ± 6.2 | 3142 ± 74 |
| LBK EN | 14.3 ± 2.5 | 13.9 ± 4.7 | 5130 ± 40 |
| Germany MN | 26.2 ± 4.4 | 55.0 ± 41.2 | 3719 ± 46 |
| Blätterhöhle MN | 18.5 ± 4.2 | 11.3 ± 4.3 | 3581 ± 86 |
| Iberia EN | 19.4 ± 2.3 | 17.5 ± 5.9 | 5107 ± 20 |
| Iberia MN | 49.8 ± 7.7 | 40.0 ± 7.0 | 3749 ± 74 |
| Iberia CA | 54.0 ± 7.1 | 58.0 ± 8.6 | 2814 ± 27 |

From Germany, we analyzed a large sample of the EN LBK culture and 11 individuals from the MN period, four of them from the Blätterhöhle site [18]. The average date of admixture for LBK (5530 ± 70 BCE) is more recent than the dates for EN/MN populations from Hungary, and the total hunter-gatherer ancestry proportion in LBK (∼4–5%) is intermediate between LBKT and ALPc. This ancestry is most closely related to a combination of KO1 and Loschbour, although the assignment of the hunter-gatherer source(s) is not statistically significant (Figure 2B; Extended Data Table 2). These results are consistent with genetic and archaeological evidence for LBK origins from the early LBKT [35], followed by additional, Central European WHG admixture after about 5500 BCE. Our “Germany MN” grouping shows increased hunter-gatherer ancestry (∼17%, most closely related to Loschbour) and a more recent average date of admixture, reflecting gene flow from hunter-gatherers after the LBK period. We successfully sequenced a total of 16 Blätterhöhle MN samples, many of them with distinct individual labels from ref. [18], although surprisingly, the genome-wide data indicated that these corresponded to only four unique individuals (Supplementary Information section 8), for which we merged libraries to increase coverage: Bla28; Bla5 (same as Bla7, Bal13, Bla26(o), Bla30, and Bla54); Bla16 (same as Bla27 and Bla59); and Bla8 (same as Bla9, Bla11, Bla24, Bla26(x), and Bla45). Based on stable isotopes, the first three of these were classified as farmers, while Bla8 had signatures associated with a hunter-gatherer-fisher lifestyle [18]. In accordance with previous results [18], we find that the group of MN farmers experienced admixture with hunter-gatherers, which we now estimate to have been at least 40%. We additionally observe admixture in the individual Bla8, with ∼25% ancestry derived from farmers. Our results thus provide evidence of asymmetric gene flow between farmers and hunter-gatherers at Blätterhöhle centered around the relatively late date of 4100 ± 120 BCE (*ALDER* dates of 10–25 generations).

In Iberia, we again see widespread evidence of local hunter-gatherer admixture, with confidently inferred LB1-related ancestry in all three population groups (EN, MN, and CA). For Iberia EN, we infer an average admixture date of 5650 ± 65 BCE, which rises to 5860 ± 110 BCE when considering only the five oldest samples (of which the earliest, CB13 [23], has an individual estimate of 5890 ± 105 BCE). Given that farming is thought to have arrived in Spain around 5500 BCE [37], these dates suggest the presence of at least a small proportion of hunter-gatherer ancestry in earlier Cardial Neolithic populations acquired along their migration route (although our admixture graph analysis only confidently detected an LB1-related component). The later Iberians have large proportions of hunter-gatherer ancestry, approximately 23% for MN (from the site of La Mina, in north-central Iberia) and 27% for CA, and also relatively old *ALDER* dates (approximately 50 generations, or 1400 years), indicating that most of the admixture occurred well before their respective sample dates. Both populations have evidence of ancestry related to LB1 and to a different WHG individual, suggesting that in contrast to the earlier admixture, the large increase in hunter-gatherer ancestry between the EN and MN had a non-local origin.

Synthesizing our time series data, we compared the observed *ALDER* dates and hunter-gatherer ancestry proportions of Neolithic populations to those estimated for simulated data under different temporal admixture scenarios (Figure 3; Extended Data Figure 4; Supplementary Information section 9). We assumed dates of 5900 BCE (Hungary) and 5500 BCE (Germany and Spain) for the onset of mixture. While none of the scenarios match the data perfectly, a reasonably good fit for Hungary is provided by a model of an initial admixture pulse (approximately 1/4 of the total hunter-gatherer ancestry observed by the end of the time series) followed by continuous gene flow (bottom solid green curve in both panels of Figure 3). Alföld and Transdanubia should be considered as separate series, but their parameters follow mostly similar trajectories, with the exception of the MN, where LBKT has a relatively old admixture date (albeit with large uncertainty) and ALPc a relatively high hunter-gatherer ancestry proportion (possibly influenced by the bias of sampling in favor of the middle and northern parts of the Alföld). Overall, even after normalizing for the different total hunter-gatherer ancestry proportions in each region, we observe a high degree of local distinctiveness, for example in the older *ALDER* dates for Iberia MN/CA and the markedly higher hunter-gatherer ancestry in Blätterhöhle. Where the simulated scenarios do not align with the real data, we can use the results to constrain the possible admixture histories in each region, e.g., by rejecting the models of a single admixture pulse or uniform continuous admixture. We also note that while the simulated data are generated under a model of gene flow from an unadmixed hunter-gatherer source population into a series of farmer populations in a single line of descent, observed admixture could also reflect immigration of new farmer populations (either via their own previous hunter-gatherer admixture or new admixture between farming populations with different proportions of hunter-gatherer ancestry). Based on archaeological evidence, such a scenario is possible, for example, for the introduction of hunter-gatherer ancestry into TDLN from Southeastern European farmers via the dispersal of the northern Balkan Vinča or Sopot cultures to Transdanubia [14,38,39].

**Figure 3.**
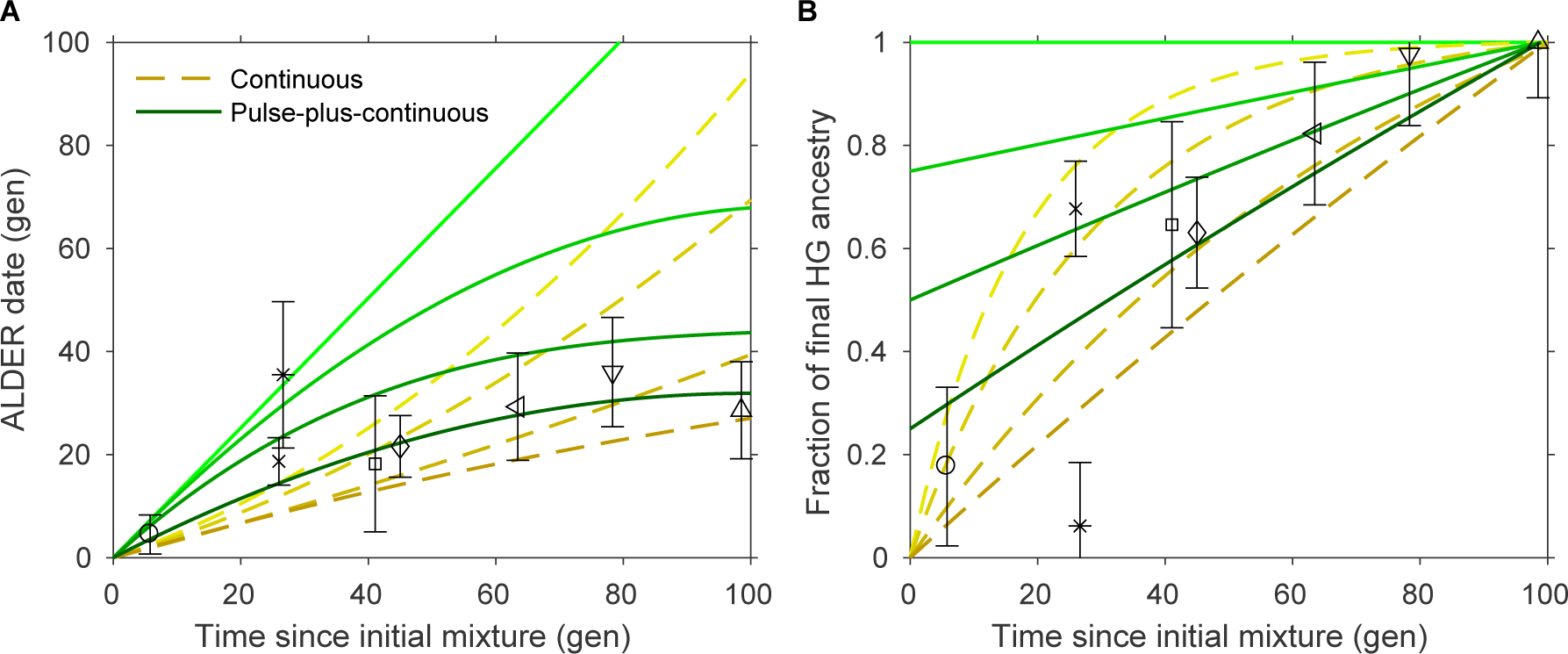
Admixture dates (A) and proportions (B) as a function of time for the Hungary time series and simulated data. Symbols are as in Figures 1 and 2, here showing population-level averages plus or minus two standard errors. Proportions and standard errors in (B) are normalized by the total hunter-gatherer ancestry in the most recent (rightmost) population in each region. Yellow dashed lines represent continuous admixture simulations: from top to bottom, diminishing 5% per generation, diminishing 3%, diminishing 1%, and uniform. Green solid lines represent pulse-plus-continuous admixture simulations: from top to bottom, all hunter-gatherer ancestry in a pulse at time zero; 3/4 of final hunter-gatherer ancestry in an initial pulse, followed by uniform continuous gene flow; 1/2 in initial pulse and the rest continuous; and 1/4 in initial pulse. Data for Germany and Iberia are shown in Extended Data Figure 4; see Supplementary Information section 9 for full details.

**Extended Data Figure 4.**
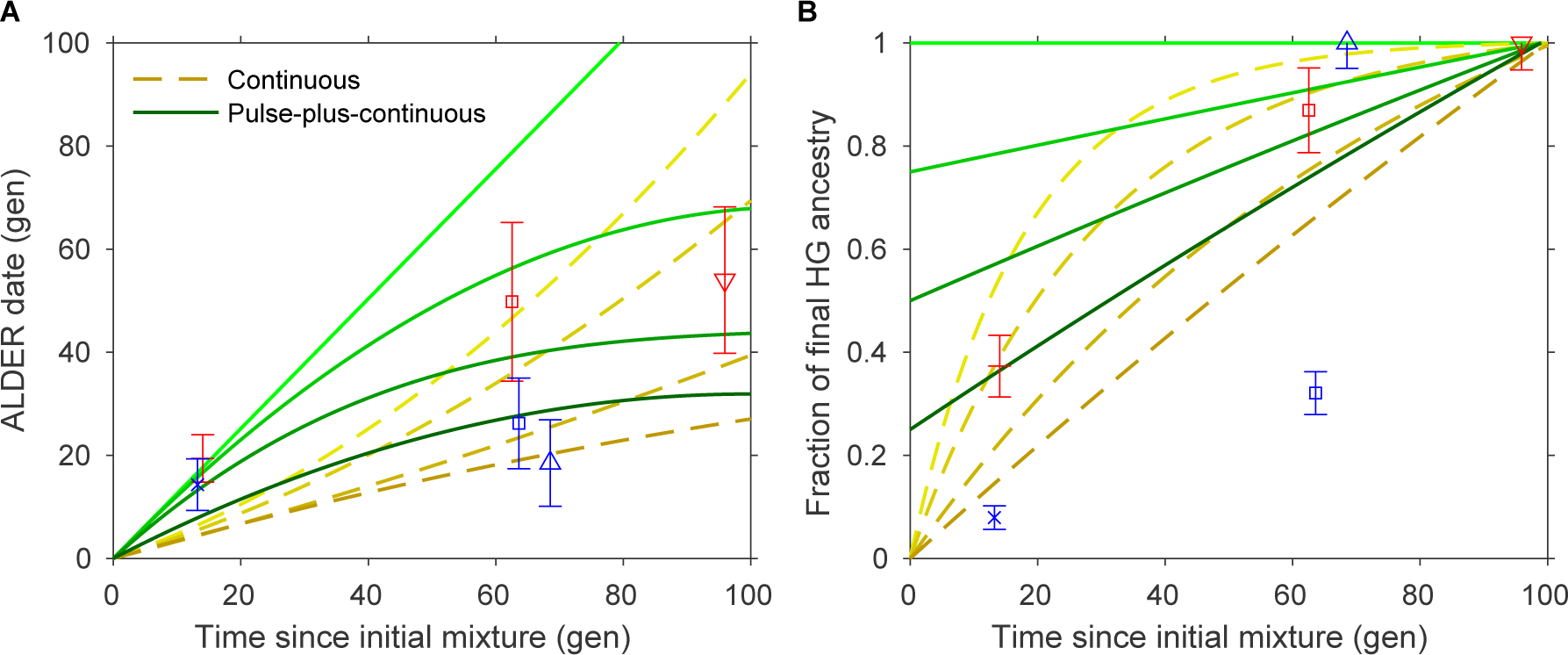
Admixture dates (A) and proportions (B) as a function of time for the Germany (blue) and Iberia (red) time series and simulated data, as in Figure 3. Symbols are as in Figures 1 and 2, here showing population-level averages plus or minus two standard errors. Proportions and standard errors in (B) are normalized by the total hunter-gatherer ancestry in the most recent (rightmost) population in each region. Yellow dashed lines represent continuous admixture simulations: from top to bottom, diminishing 5% per generation, diminishing 3%, diminishing 1%, and uniform. Green solid lines represent pulse-plus-continuous admixture simulations: from top to bottom, all hunter-gatherer ancestry in a pulse at time zero; 3/4 of final hunter-gatherer ancestry in an initial pulse, followed by uniform continuous gene flow; 1/2 in initial pulse and the rest continuous; and 1/4 in initial pulse.

Our results provide greatly increased detail in understanding population interactions and admixture during the European Neolithic. In each of our three study regions, the arrival of farmers prompted admixture with local hunter-gatherers which unfolded over thousands of years: almost all sampled populations from subsequent time periods have more hunter-gatherer ancestry and more recent dates of admixture than their local predecessors, suggesting recurrent changes in genetic composition and significant hunter-gatherer gene flow beyond initial contact. These transformations left distinct signatures in each region, implying that they likely resulted from a complex web of local interactions rather than a uniform demographic phenomenon. Our transect of Hungary, in particular, with representative samples from many archaeological cultures across the region and throughout the Neolithic and Chalcolithic, illustrates the power of dense ancient DNA time series. Future work with similar data sets and statistical models promises to teach us much more about population transformations in space and time.

## Methods

### Experimental procedures

Prehistoric teeth and petrous bone samples from Hungary were taken under sterile conditions in the Hungarian Museums and anthropological collections. Sixty-six samples were documented, cleaned, and ground into powder in the Anthropological Department of the Johannes Gutenberg University of Mainz, during the course of the German Research Foundation project AL 287-10-1, and the other 20 were prepared in Budapest, in the Laboratory of Archaeogenetics of the Institute of Archaeology, Research Centre for the Humanities, Hungarian Academy of Sciences, following published protocols [35]. DNA was extracted in Budapest using 0.08–0.11g powder via published methods [40], using High Pure Viral NA Large Volume Kit columns (Roche) [21, 41]. DNA extractions were tested by PCR, amplifying the 16117–16233 bp fragment of the mitochondrial genome, and visualized on a 2% agarose gel. DNA libraries were prepared from clean and successful extraction batches using UDG-half and no-UDG treated methods [10, 42]. We included milling (hydroxylapatite blanks to control for cleanness) and extraction negative controls in every batch. Barcode adapter ligated libraries were amplified with TwistAmp Basic (Twist DX Ltd), purified with Agencourt AMPure XP (Beckman Coulter), and checked on 3% agarose gel [10]. Library concentration was measured on a Qubit 2.0 fluorometer. Promising libraries after initial quality control analysis were shipped to Harvard Medical School, where further processing took place. All other samples were prepared similarly in dedicated clean rooms at Harvard Medical School and the University of Adelaide in accordance with published methods [10,12,21].

We initially screened the libraries by examining mitochondrial DNA (mtDNA) content via in-solution hybridization to a set of mtDNA probes [43], using a protocol described previously [10, 21]. Libraries with good screening results—limited evidence of contamination, reasonable damage profiles, and substantial coverage on targeted segments—were enriched for a genome-wide set of ∼1.2 million SNPs [12, 21] and sequenced to greater depth. Raw sequencing data were processed by trimming bar codes and adapters, merging read pairs with at least 15 base pairs of overlapping sequence, and mapping to the human reference genome (version hg19). Reads were filtered for mapping and base quality, duplicate molecules were removed, and two terminal bases were clipped to eliminate damage (five for UDG-minus libraries) [10]. All libraries had a rate of at least 4.8% C-to-T substitutions in the final base of screening sequencing reads (Supplementary Table 1), consistent with damage patterns expected for authentic ancient DNA [42, 44]. Pseudohaploid genotypes at each SNP were called by choosing one allele at random from among mapped reads.

Mitochondrial DNA sequences were reassembled in Geneious R10 to rCRS [45] and RSRS [46], and SNPs with at least 3x coverage and a minimum variant frequency of 0.7 were called. The assembly and the resulting list of SNPs were double-checked against phylotree.org (mtDNA tree Build 17; 18 Feb 2016). Haplotype calls are given in Table SXX (in progress). On the Y chromosome, 15,100 SNPs were targeted and sequenced, and the detected derived and ancestral alleles were compared to the ISOGG Y-tree (www.isogg.org) version 12.34, updated on 5th February 2017. Haplogroup definitions are detailed in Supplementary Information section 3.

We merged libraries from the same individual (for those with more than one) and then combined our new samples with genome-wide data from the literature (ancient individuals as described and as listed in Extended Data Table 1 and present-day individuals from the SGDP [47]) using all autosomal SNPs (∼1.15 million) from our target set. For two replications of our admixture graph analyses, we restricted either to the subset of transversions (∼280K SNPs) or to the subset from panels 4 and 5 of the Affymetrix Human Origins array (ascertained as heterozygous in a San or Yoruba individual; ∼260K SNPs). For PCA (Extended Data Figure 1), we merged with a large set of present-day samples [21] and used all autosomal Human Origins SNPs (∼593K).

To test for possible contamination, we used contamMix [48] and ANGSD [49] to estimate rates of apparent heterozygosity in haploid genome regions (mtDNA and the X chromosome in males, respectively). Any samples with > 5% mtDNA mismatching or > 2% X contamination were excluded from further analyses, with the exception of Blätterhöhle individual Bla5, which was differentiated from all other samples with signs of mtDNA contamination by its substantially higher coverage on the nuclear genome (∼5x, all others < 1x, median ∼0.1x; Supplementary Table 1), was comprised of data from six merged libraries, and did not deviate noticeably from the other Blätterhöhle individuals in our analyses. We also removed samples identified as clear outliers in PCA or with significant population genetic differences between all sequencing data and genotypes called only from sequences displaying ancient DNA damage signatures. A total of 17 samples were excluded based on one of these criteria. For individual-level f-statistic analyses (Figure 2A–B), we restricted to samples with a maximum level of uncertainty, defined as a standard error of at most 7×10^−4^ for the statistic *f*_4_(Mbuti, WHG; Anatolia, *X*). This threshold (corresponding to an average coverage of approximately 0.05, or ∼60K SNPs hit at least once) was met by 87 of the 110 samples passing QC (and 49 of the 50 samples from the literature). We did not impose such a threshold for *ALDER* analyses, but because low coverage results in a weaker signal, only one of the 23 high-uncertainty individuals in our primary data set provided an *ALDER* date (as compared to 86 of the 128 low-uncertainty individuals).

### Population assignments for analyses

In most cases, population groupings were used that correspond to archaeological culture assignments based on chronology, geography, and material culture traits. Occasionally, we merged populations that appeared similar genetically in order to increase power: we pooled samples from all phases and groups of the eastern Hungarian MN into a single ALPc population; merged six Sopot with eight Lengyel individuals for the western Hungarian TDLN; combined one Hunyadihalom (Middle CA from the Danube-Tisza interfluve in central Hungary) with Lasinja; pooled four LBK samples from Stuttgart with the majority from farther to the northeast (primarily Halberstadt); and merged several cultures of the German MN into a single group. Other populations vary in their degrees of date and site heterogeneity, with Iberia MN the most homogeneous and Iberia EN and CA among the least (Extended Data Table 1; Supplementary Table 1). For our main analyses, we excluded the Vinča and Tiszapolgár population groups because they lacked sufficient high-quality data.

We note that the designations EN, MN, LN, and CA have different meanings in different areas. For our study regions, each term generally refers to an earlier period in Hungary than in Germany and Spain (for example, ALPc and LBKT MN in Hungary are roughly contemporaneous with LBK and Iberia EN). In order to maintain agreement with the archaeological literature, we use the established definitions, with the appropriate word of caution that they should be treated separately in each region.

### Sample dates

We report 44 newly obtained accelerator mass spectrometry (AMS) radiocarbon dates for Neolithic individuals (38 direct, 6 indirect), focusing on representative high-quality samples from each site and any samples with chronological uncertainty. These are combined with 57 radiocarbon dates from the literature [9, 10, 12, 22, 23, 35, 38, 39, 50]. We report the 95.4% calibrated confidence intervals (CI) from OxCal [51] version 4.2 with the IntCal13 calibration curve [52] in Extended Data Table 1. For use in *ALDER* analyses (Supplementary Information section 7), we use the mean and standard deviation of the calibrated date distributions; while the distributions are non-normal, we find that on average the mean plus or minus two standard deviations contains more than 95.4% of the probability density. For samples without direct radiocarbon dates but with dates from other samples or materials at the same site, we form a conservative 95.4% CI by taking the minimum and maximum bounds of any of the calibrated CIs from the site. Finally, for the remaining samples, we use plausible date ranges based on archaeological context; we assume independence across individuals but as a result take a conservative approach and treat the assigned range as ± one standard error (e.g., an estimated range of 4800–4500 BCE becomes 4650 ± 150 BCE).

## Acknowledgments

We thank Iosif Lazaridis, Po-Ru Loh, Iain Mathieson, Iñigo Olalde, Nick Patterson, and Pontus Skoglund for helpful comments and suggestions; Johannes Krause for providing the Stuttgart sample for which we generated a new library in this study; and Bálint Havasi (Balaton Museum), György V. Székely (Katona József Museum), Csilla Farkas (Dobó István Museum), Borbála Nagy (Herman Ottó Museum), I. Pap, A. Kustár, T. Hajdu (Hungarian Natural History Museum), J. Odor (Wosinsky Mór Museum), E. Nagy (Janus Pannonius Museum), P. Rácz (King St. Stephen Museum), L. Szathmáry (Debrecen University), N. Kalicz, V. Voicsek, O. Vajda-Kiss, V. Majerik, and I. Kővári for assistance with samples. This work was supported by the Australian Research Council (grant DP130102158; B.L. and W.H.), Hungarian National Research, Development and Innovation Office (K 119540; B.M.), German Research Foundation (Al 287 / 7-1, 10-1, 14-1; K.W.A.), FEDER and Ministry of Economy and Competitiveness of Spain (BFU201564699-P; C.L.-F.), National Science Foundation (HOMINID grant BCS-1032255; D.R.), National Institutes of Health (NIGMS grant GM100233; D.R.), and Howard Hughes Medical Institute (D.R.).

## Author contributions

A.S.-N., J.B., E.B., K.W.A., C.L.-F., W.H., and D.R. designed and supervised the study. B.G.M., K.K., K.O., M.B., T.M., A.O., J.J., T.P., F.H., P.C., J.K., K.Se., A.A., P.R., J.R., J.P.B., S.F., G.S., Z.T., E.G.N., J.D., E.M., G.P., L.M., B.M., Z.B., J.F.-E., J.A.M.A., C.A.F., J.J.E., R.B., J.O., K.Sc., H.M., A.C., J.B., E.B., K.W.A., C.L.-F., and W.H. provided samples and assembled archaeological and anthropological information. A.S.-N., A.P., B.S., V.K., N.R., K.St., M.F., M.M., J.O., N.B., E.H., S.N., and B.L. performed laboratory work. M.L., A.S.-N., S.M., and D.R. analyzed genetic data. M.L., A.S.-N., and D.R. wrote the manuscript with input from all coauthors.

